# Modeling bi-modality improves characterization of cell cycle on gene expression in single cells

**DOI:** 10.1101/002295

**Authors:** Andrew McDavid, Lucas Dennis, Patrick Danaher, Greg Finak, Michael Krouse, Alice Wang, Philippa Webster, Joseph Beechem, Raphael Gottardo

**Affiliations:** Department of Statistics, University of Washington, Seattle, WA 98195, USA; Vaccine and Infectious Disease Division, Fred Hutchinson Cancer Research Center, Seattle, WA 98109, USA; NanoString Technologies, Seattle, WA 98118, USA; BD Biosciences, San Jose CA 95131, USA

## Abstract

Advances in high-throughput, single cell gene expression are allowing interrogation of cell heterogeneity. However, there is concern that the cell cycle phase of a cell might bias characterizations of gene expression at the single-cell level. We assess the effect of cell cycle phase on gene expression in single cells by measuring 333 genes in 930 cells across three phases and three cell lines. We determine each cell’s phase non-invasively without chemical arrest and use it as a covariate in tests of differential expression. We observe bi-modal gene expression, a previously-described phenomenon, wherein the expression of otherwise abundant genes is either strongly positive, or undetectable within individual cells. This bi-modality is likely both biologically and technically driven. Irrespective of its source, we show that it should be modeled to draw accurate inferences from single cell expression experiments. To this end, we propose a semi-continuous modeling framework based on the generalized linear model, and use it to characterize genes with consistent cell cycle effects across three cell lines. Our new computational framework improves the detection of previously characterized cell-cycle genes compared to approaches that do not account for the bi-modality of single-cell data. We use our semi-continuous modelling framework to estimate single cell gene co-expression networks. These networks suggest that in addition to having phase-dependent shifts in expression (when averaged over many cells), some, but not all, canonical cell cycle genes tend to be co-expressed in groups in single cells. We estimate the amount of single cell expression variability attributable to the cell cycle. We find that the cell cycle explains only 5%-17% of expression variability, suggesting that the cell cycle will not tend to be a large nuisance factor in analysis of the single cell transcriptome.

## Introduction

With the advent of single cell expression profiling [1–4], the assessment of cell population heterogeneity and identification of cell subpopulations from mRNA expression is achievable [5–7]. However, at the single cell level, there is concern that cell cycle might interfere with the characterization of gene expression variability [8]. As many biological samples are prepared from asynchronous cell populations, where each cell is in an unknown phase of the cell cycle, it is imperative to understand the impact of cell cycle in order to account for its effect on observed expression patterns and downstream data analysis. Here, we have measured mRNA expression and cell cycle from 930 single cells derived from three cell lines in order to explore this hypothesis.

A distinctive feature of single-cell gene expression data is the bimodality of expression values. Genes can be on (and a positive expression measure is recorded) or off (and the recorded expression is zero or negligible)[9,10]. This dichotomous characteristic of the data prevents use of the typical tools of designed experiments such as linear modeling and analysis of variance (ANOVA). We develop a novel computational framework to overcome this problem. First, a probabilistic mixture model-based framework allows the separation of positive expression values from background noise using gene-specific thresholds. After signal separation by thresholding, we model separately the frequency of expression (the fraction of cells expressing a gene) and the continuous, positive expression values. Our semi-continuous framework combines evidence from the two salient parameters of single cell expression in a statistically appropriate manner, an approach dubbed the Hurdle model[11,12]. Extending our previous proposal of a two-sample semi-continuous test akin to the two-sample *t-test*, our new framework allows for testing arbitrary contrasts and allows the use of variance components/mixed models, thus bringing to bear the full power of the general linear model.

The Hurdle model allows us to identify many genes with an archetypal cell cycle expression pattern despite a frequently bimodal distribution of expression. It also suggests that stochastic variation in single cell gene expression is relatively large compared to the effect of cell cycle. We find that even in the most tightly regulated gene, cell cycle explains only 27% of the variability, while in the median gene in our data set, cell cycle explains 5%-18% of the variability, depending on the assumptions we make regarding latent technical variability. The semi-continuous model also provides a framework for estimating co-expression networks - in which edges connect genes whose partial correlations remain after removing the effect of all other genes - while adjusting for population-level nuisance factors that could bias network inference. Applying this framework to our data, we show that only a subset of canonical cell cycle genes are highly co-expressed in single cells.

## Results

### Periodic expression associated with cell cycle is observed at the single-cell level

In order to assess differential expression associated with actively cycling cells, expression of 333 genes was interrogated in 930 cells, across three cell lines: H9 (HTB-176), MDA-MB-231 (HTB-26), and PC3 (CRL-1435) (Figure 1A). Single cell expression was measured from flow-sorted cells and compared between cell cycle phases and cell lines via nCounter single cell profiling, a multiplexed hybridization-based detection technology that utilizes fluorescent barcodes to count individual target nucleic acid molecules [13]. This platform has been recently adapted to enable expression profiling from single cells via hybridization after a multiplexed target enrichment (MTE) in which mRNA is first converted to cDNA and then amplified [14].

**Figure 1.**
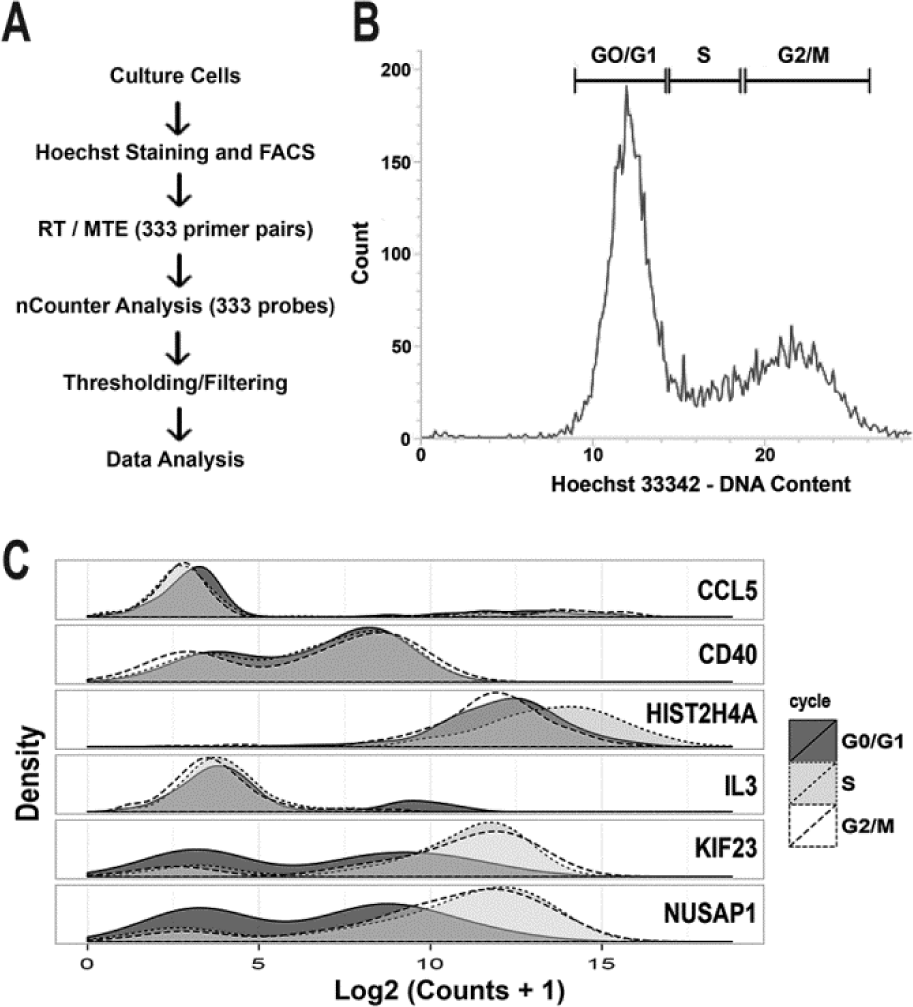
Individual cells were flow sorted by DNA content, and gene expression profiled. (A) H9, MB231, and PC3 cells were cultured and sorted into lysis buffer. The resulting lysate was amplified via multiplexed target enrichment (MTE) and digital counts of expression were optically read via nCounter. (B) Individual cells were sorted into three populations based on retention of Hoechst dye (G0/G1, S and G2/M). (C) The density distribution of log counts for each gene was generally bimodal with some genes showing clear changes in distribution between cell cycle phase.

Each cell was categorized as being in G0/G1, S or G2/M phase by measuring DNA content via flow-cytometry based on retention of Hoechst dye (Figure 1B and S1)[15]. Probes were selected for cell cycle associated genes (n = 119). These genes provided coverage of the entire cell cycle (Data Set S1) based on peak expression and periodicity information obtained from Cyclebase, an integrated database of bulk cell cycle expression profiling experiments that scores and ranks genes based on strength of evidence for a cell cycle associated expression pattern[16]. Probes were also included for non-cell-cycle associated genes with primary roles in the inflammatory response, and housekeeping controls without a Cyclebase ranking (n=214). We denote probes with a Cyclebase rank (i.e. genes with the strongest evidence for cell cycle associated periodic expression) as the *ranked* set.

253 genes were expressed and passed quality control (see Methods). Genes showed a bimodal expression pattern in log-transformed mRNA levels (Figure 2), consistent with a burst-model of “on/off” transcription at the single cell level [17] and consistent with the kinetics of PCR amplification with low starting template concentrations, described by us and other authors [9,10].

**Figure 2.**
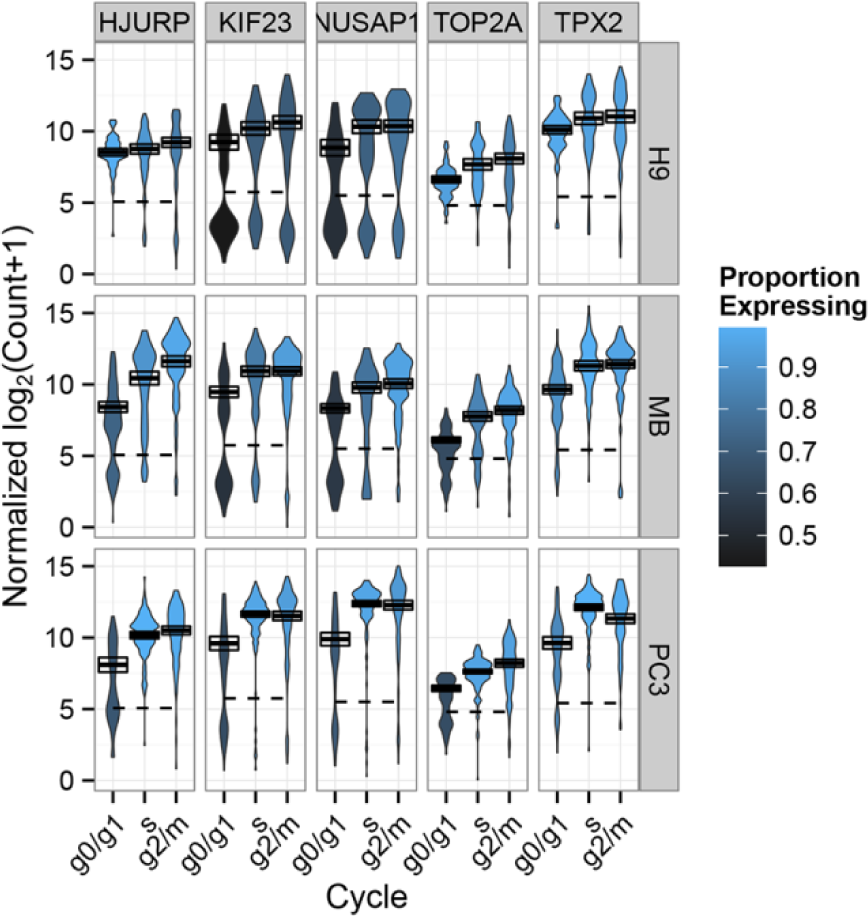
Top 5 cell cycle genes detected by the Hurdle model. A violin plot shows the density of log counts of mRNA in each condition. The expression threshold estimated for each gene is shown as a dashed line, so that the ratio of area above the dashed line reflects the proportion of cells expressing a gene. Blue shades of the violin depict genes with more expressing cells in a condition. The positive mean and 95% confidence interval is depicted as a box with solid line. The Hurdle model (see Methods) combines evidence for changes in either of these parameters, after adjusting for cell line, to determine statistical significance.

Expression levels for each gene were most different between cell lines (Figure 2). Many genes, including those in the *ranked* set showed cell line-specific expression patterns. For example, expression of TOP2A in G0/G1 varied from 70% of cells in MB-231 and PC3 to nearly universal in H9. This cell line effect was a nuisance factor we needed to adjust for in differential expression tests on cell cycle.

Nonetheless, many genes from the *ranked* set, such as KIF23, TOP2A, HJURP, NUSAP1, and TPX2 exhibited expression patterns consistent with cell cycle regulation (Figure 2). Figure 2 also reveals that changes in both the positive expression mean (i.e. the mean over the cells expressing that gene; PEM), and changes in the frequency of cells expressing a gene, occur throughout the cell cycle. The frequency and PEM in these genes also vary widely between cell lines, so it was important to adjust for cell line effects for accurate assessment of differential expression.

In order to test for significant differences in expression between cell cycle phases that were consistent across cell lines, we developed an ANOVA-like model (Hurdle model, see Methods) that permits adjustment for additive effects due to cell line. The Hurdle model improves the power to detect changes in single-cell expression by testing both the frequency of expression (corresponding to the relative distribution of cells between the two modes), and the PEM. Combining evidence from the discrete and continuous components of the data provides better sensitivity to changes in expression compared to test statistics based on frequencies of expression (discrete) or on the PEM (continuous) alone; or a union test (see Materials and Methods) while remaining competitive in specificity (Figures S3, S4)

Within the three cell lines tested here, *significant* differential expression (Bonferroni-adjusted for 253 tests at P<0.05) was observed for 78 genes in the *ranked* set and 28 genes in the *unranked* set (Figure 3A). Genes showing the strongest cell cycle associated expression patterns in bulk measurements were more likely to be identified as significant in the single-cell populations (Figure 3A-B).

**Figure 3.**
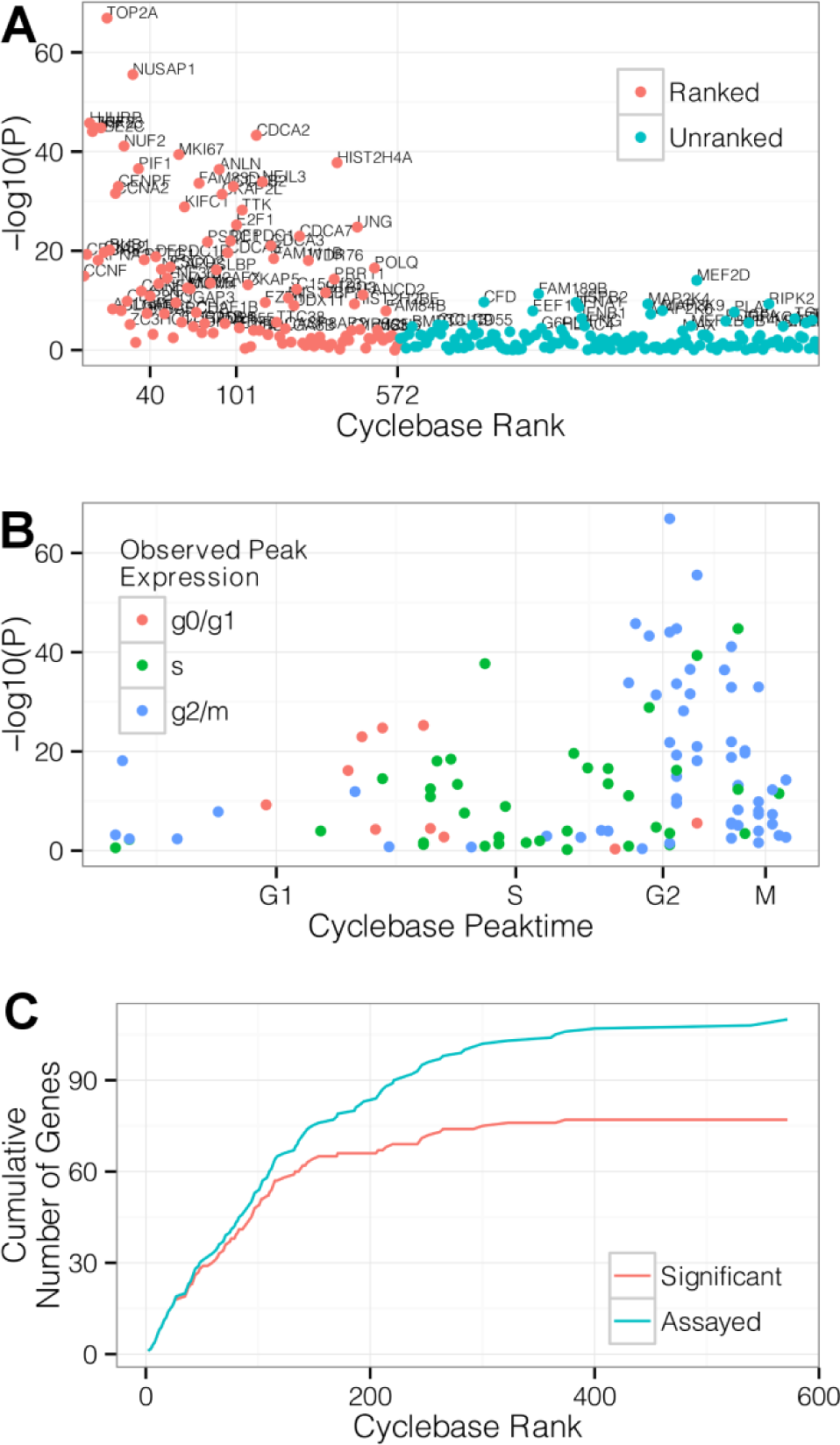
The Hurdle model identifies genes with cell cycle phase-dependent expression. (A) Hurdle model strength of evidence of cell cycle phase dependent expression for genes within our panel versus Cyclebase rank. P-values (-log10(p)) are shown on the y-axis. *Ranked* genes (red) are ordered on the x-axis according to Cyclebase rank. Unranked genes (blue) appear in alphabetical order. Genes significant after Bonferroni adjustment are annotated with their names (B) Hurdle model strength of evidence of cell cycle phase dependent expression in *ranked* genes versus phase of peak expression estimated from bulk data in Cyclebase. Experimentally observed peak times broadly match the times estimated from bulk data. Concordance in observed peak times is greater for genes with stronger evidence of differential expression. (C) Cumulative number of significant (red) or all (blue) *ranked* genes versus Cyclebase rank. Genes with lower Cyclebase rank, and hence stronger evidence of cycle regulation in bulk expression, are detected more often than genes with weaker evidence as shown by the minimal gap between significant (red) and all (blue) *ranked* gene lines at Cyclebase rank <150.

For each gene, peak time was determined based on the phase (G0/G1, S or G2/M) with maximum average expression across all cell lines. Despite large cell-line-specific expression variability, peak times were broadly consistent with Cyclebase annotations (Figure 3C), and especially so within the subset of genes with strongest evidence of cycle regulation in our data (e.g. Bonferroni significant at P < 0.05).

The majority of genes in the *unranked* set (115/143 or 80%) did not exhibit significant cell cycle effects, in concordance with their primary roles in functions unrelated to the cell cycle. Of the 28 *unranked* genes that exhibited a significant cell cycle phase association, we noted genes involved in cytoskeletal organization (PLAT), proliferation (PDGFA), and signaling pathways (IFNA1, IFNB1) that have been previously demonstrated to modulate progression through the cell cycle [18].

### Cell cycle explains a small portion of the gene expression variability

It has been argued that a substantial portion of the stochastic variability observed in single cell gene expression experiments may be caused by global changes in transcription due to cell cycling [19]. We explore this idea by examining the proportional change in the Hurdle model fit associated with inclusion and omission of cell cycle as an explanatory variable. Because the Hurdle model accounts for both the dichotomous (on/off) and continuous nature of single cell data, the change in deviance (generalized linear model log-likelihood) between nested models can be used to calculate the amount of variability explained by cell cycle. The total deviance can be partitioned into components corresponding to cell cycle effects, nuisance effects described below, and residual effects. The ratio of cell cycle deviance to the sum of cell cycle plus residual deviance can then be interpreted as the analog to the coefficient of determination in linear least squares.

We consider expression changes due to main effects and interactions of *cell cycle* by *cell line* and account for amplification efficiency and average cell line effect (see Materials and Methods). Only modest amounts of the single cell expression variability can be explained by cell cycle (Figure 4). Within the *ranked* gene set, cell cycle phase explains 8% of the deviance in the median gene and 27% of the deviance in the top gene (TOP2A). In *unranked* genes, phase explains only 5% of the deviance in the median gene.

**Figure 4.**
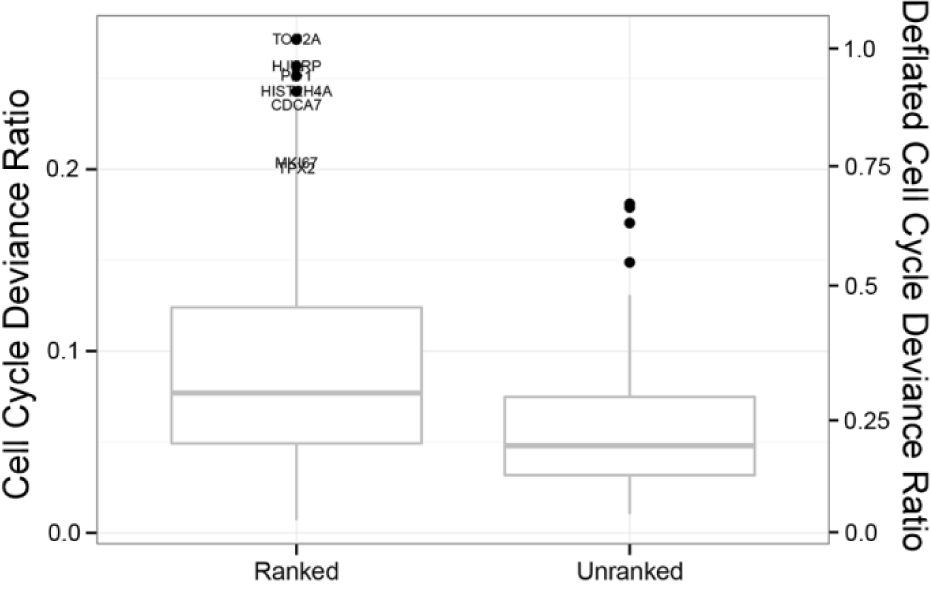
Box and Whiskers Plot of Cell Cycle Deviance Ratio in Ranked and Unranked Genes. The proportion of stochastic variability in the Hurdle Model explained by cell cycle is shown on the primary y-axis (left) for *ranked* and *unranked* genes, with box giving the 25th, 50th and 75th percentiles, and whiskers showing 1.5 times the inter-quartile range. The deflated scale on the secondary y-axis (right) shows the deviance as a percentage of the most completely explained gene (TOP2A, 27%) and is intended as an upper bound for the amount of remaining biological deviance in non-cell-cycle genes. Under this conservative rescaling, cell cycle explains only 25% of the deviance in 75% of unranked genes.

To derive these estimates, it is important to be able to account for the nuisance factors by using the Hurdle model. If cell-to-cell variation in amplification efficiency is not removed, we underestimate the explanatory power of cell cycle on in the median ranked gene by 26% since the unmodeled deviance would include this large additional component. Similarly, other unmeasured factors may inflate the residual deviance and attenuate the apparent role of cell cycle. These factors could include errors in inferring the cell cycle phase via FACS or imperfect modeling of changes in amplification or detection efficiency between samples. To guard against this attenuation, we set an upper bound on cell-cycle-dependent variation as follows: We suppose that transcription of the gene with the most deviance attributable to cell cycle (TOP2A, 27%) would be entirely regulated in a phase-dependent manner, and we characterize other genes’ cell-cycle-dependent deviance relative to this maximum. For example, a gene with 13.5% cell-cycle-dependent deviance has half as strong a cell cycle effect as TOP2A, leading to the conclusion that at most 50% of this gene’s deviance could be attributable to cell cycle. Even under these generous upper bounds, cell cycle phase explains only 18% (eg, .05/.27) and 29% (eg,.08/.27) of the deviance in the median gene in the *unranked* and *ranked* sets, respectively, suggesting that even when allowing for cell line-specific cell cycle effects, cycle is generally a small factor, compared to residual variability, in gene expression variability in the human transcriptome.

### Network analysis reveals gene co-expression at the single-cell level

Single-cell gene expression data sets have the resolution to reveal not only differential expression in response to biological variables like cell cycle phase, but also to provide insight into co-expression between genes at the cellular level (e.g. the influence of one gene on another’s expression or the sharing of upstream regulatory elements). In bulk-gene expression data (e.g. microarrays), apparent co-expression arises from tissue-level factors inducing shared marginal changes in genes. For example, different radiation doses in samples will induce correlation amongst all the genes affected by radiation, regardless of whether these genes interact or even participate in the same biological processes. In contrast, single cell data allow isolation of co-expression arising from cellular-level factors, giving access to more fundamental biological relationships. If two genes are correlated across cells drawn from the same environment, then the two genes are likely to share an intimate biological relationship: they may be regulated by the same transcription factor, or one gene may directly regulate the other. The distinction between cellular and marginal co-expression follows from a probabilistic identity on conditional covariances (see Materials and Methods).

When cell cycle is not adjusted for (Figure 5 D-F), known cell cycle genes with strong evidence of marginal regulation comprise the majority of the network. These genes generally peak in phase G2/M, suggesting that the co-expression is mostly driven by the coincident peak in average expression. The networks adjusted for cell cycle at least partially remove marginal effects (Figure 5 A-C). In some cell cycle genes, substantial evidence for co-expression remains, but now additional co-expression is detected in genes without a previously described cell cycle role. In the unadjusted estimates, marginal shifts in expression in canonical cell cycle genes overwhelm subtler co-expression in unranked genes. Even though cell cycle variability is modest compared to residual variability, cell cycle is a substantial source of *biological* variability in the ranked genes and is in a sense confounded with the co-expression patterns.

**Figure 5.**
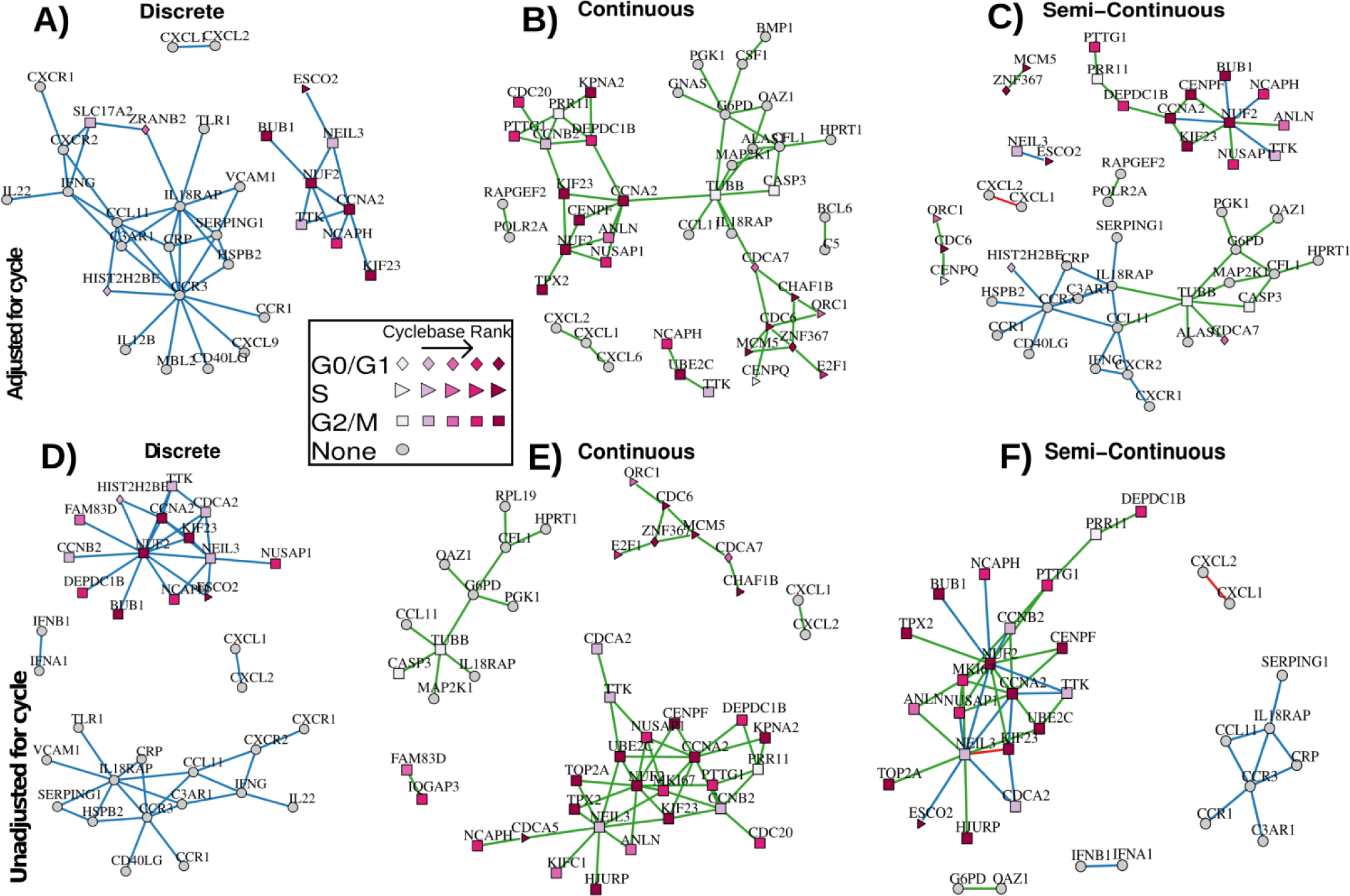
Coexpression networks estimated using the Hurdle Model. Data from three cell lines and three cycles are combined and adjusted for additive effects of cell line and pre-amplification efficiency. Networks of the top 60 edges (ranked by partial correlation) using logistic regressions on discretized expression (A,D), linear regressions on positive, continuous expression values (B,E), and combining the top 30 edges from discrete and continuous components are shown (C,F). Panels A-C adjust for additive cell cycle effects, while panels D-F are unadjusted. The shape of the glyph corresponds to the cycle with peak expression from cyclebase, while the saturation of the glyph corresponds to the ranking. Blue and green edges are partial correlations detected from discrete expression and continuous expression, respectively. Red edges are detected in both discrete and continuous expression.

In an attempt to quantify the performance of the Hurdle model and the effect of cell-cycle adjustments, we examined network properties when varying the number of edges. We call an edge *peaktime concordant* if it connects nodes that have the same peaktime annotated in cycle base (eg, G0/G1-G0/G1 or S-S). Over a range of network densities (30-240 edges) the unadjusted Hurdle or Raw networks contain between 45%-80% peaktime concordant edges, while the adjusted Hurdle contains only 32%–38% peaktime concordant edges.

Cell cycle adjustment in networks estimated on the raw data is not very effective compared to the unadjusted, raw networks (Figure S6). This is unsurprising, as this would occur when the model for the mean of the response is mis-specified, as is true when ignoring the bi-modality that the data exhibit (eg, Figures 2 and S2). If the Hurdle model is correct and cell cycle is additive, then the identity link cannot recover this additivity. On the other hand, the Hurdle model can still recover an additive mean model under a linear link by taking the discrete coefficient estimates to be null. Overall, the adjusted and unadjusted Hurdle networks in Figure 5 are rather different, sharing 39% of nodes (Jaccard similarity) and 51% of edges (Hamming Distance/#edges).

Combining both discrete and continuous networks (with the top 30 edges from discrete and continuous networks) allows a richer set of genes to be characterized. When discrete expression is used alone, networks primarily consist of G2/M peaking genes and unranked genes (Figure 5A). When positive, continuous expression is also used, S and G0/G1 peaking genes enter the networks (Figure 5B-C).

The adjusted, semi-continuous network depicted in Figure 5C consists of two primary sub-networks, one consisting entirely of ranked genes, and another largely consisting of weakly ranked and unranked genes. While we cannot rule out that measurement error of the inferred cycle is not partially responsible for the persistence of a subset of ranked genes, previously described mutual regulation in RNA-interference experiments [20] of some of these genes suggests that this subset is co-expressed at the single cell level as opposed to being co-expressed on average at the population level. The sub-network of ranked genes contains the central node of NUF2, a highly-conserved protein required for stable kinetochore localization of centromere-associated protein E (CENP-E) [21]. NUF2 is connected to other actors in mitotic organization such as ANLN, KIF23, and CENPF, as well as the check-point genes CCNA2 and BUB1, reflecting the central role of these genes in mitosis.

The sub-network of primarily unranked genes contains two key nodes: TUBB and CCR3. The predominance of genes associated with cell growth, like TUBB, and transmembrane proteins, like CCR3, in the unranked cluster is likely related to the actively dividing nature of the profiled cells, i.e. dividing cells must generate new scaffolding and membrane-related materials to support growth. This relatively large sub-network of unranked and weakly ranked genes is largely missed by the unadjusted analysis that is biased by the population level cell-cycle effect.

## Discussion

Stochastic, bimodal expression is a hallmark of single cell data [22–24]. Within a population of cells, detectable expression for any given gene typically resides in one of two modes, corresponding to an “on” or “off” state. Both technical and biological factors likely contribute to this bimodality. Quantities of some species of cDNA may be minute after reverse-transcription, and in this case random variation in the number of template-primer-enzyme complexes that form during each annealing phase may dominate the kinetics of the PCR [25]. But regardless of its origin, modeling bimodality improves the power of differential expression tests.

Here, we show how the Hurdle model can be adapted to complex study designs, extending our previous results describing its use for two-sample comparisons. We demonstrate the model’s ability to identify many genes with a periodic expression pattern from asynchronously cultured cells utilizing a combination of FACS sorting and these new analytical techniques, including genes with little previous evidence of cell cycle associated periodic expression like MEF2D [26] and FAM189B. The Hurdle model is able to identify phase-dependent patterns of expression despite the fact that G2 and M phases are indistinguishable by DNA content. The similar rank ordering of differentially expressed genes in our single cell experiment as compared to bulk experiments and concordance in the phase of peak expression demonstrates the power of the Hurdle model. While we have applied the Hurdle model to our specific problem, the approach is general and can be applied to test any effect of interest in a single-cell gene expression dataset. We offer this modeling framework as an R package for other interested users at github.com/RGLab/SingleCellAssay.

Although we recommend the Hurdle model in general for testing for differential expression, it should be noted that its desirability is contingent on the frequency of the gene under consideration. For example, if a gene is highly expressed (eg, > 90% expression), then the information to be derived from the 10% of cells that do not express a gene may not be worth the cost of an extra degree of freedom in the chi-square null distribution of the test statistic. However, even when this is the case, the Hurdle model might be preferred for methodological simplicity, since it is powered—although perhaps not always optimally—regardless of expression frequency, and does not require extensive pre-test simulations of power to yield acceptable performance. The data set considered here offers a relatively stringent test of the relative sensitivity of the Hurdle model, owing to the high expression frequency of the genes in this experiment (interquartile range ranked genes: .7-.9; unranked genes: .56-.88).

Single cell data also allows unparalleled resolution of genes’ co-expression patterns. While bulk expression data can reveal correlation induced by varying biological conditions, single-cell data has the possibility to reveal co-expression driven by shared regulatory elements within the cell. However, when inferring gene expression networks, it is important to adjust for population level covariates that could confound the network estimation, especially for genes that are marginally affected by such a population level covariate (like known cell cycle genes in our experiment.) By measuring a limited set of cell cycle associated genes, we are able to identify a network of co-expressed genes with known roles in cell cycle regulation even after adjusting for cell cycle phase. It should be noted that the unadjusted network estimate would be appropriate in some circumstances, for example when a summary of the co-expression occurring on *average* in the *population* of cells is desired, as opposed to inference of co-expression occurring conditionally within defined subsets.

Work remains to derive network estimators that optimally combine information from discrete and continuous portions. Our current approach is likely theoretically naïve, since it is essentially a union test of the discrete and continuous portions, rather than a summation of signal from the two domains. We also have left unresolved the asymptotic consistency of our proposed network procedure under dimensional scaling.

It is crucial to understand the relationship between cell cycle and the stochastic nature of single cell expression as it determines the magnitude of the cell cycle’s distorting effect on single cell analyses. In contrast to earlier estimates of Zopf *et al*. [19] we find little evidence of periodic regulation of expression among non-cell cycle associated genes. Our results are consistent with genome-wide mRNA profiling efforts utilizing bulk expression methodologies in mammalian cells where genes with cycle-dependent periodic expression patterns are limited and well-characterized [16,27,28]. Disparity between our findings and those of Zopf *et al*. may arise from differences between yeast and mammalian cells. Moreover, Zopf *et al.* primarily focus on a single, synthetic promoter while we sample hundreds of transcripts presumably driven by many different promoters. Whether the substantial remaining variability is inherent to the human single cell, or due to thus far latent, unmeasured biological variables remains to be explored.

## Materials and Methods

### Cell Lines and Flow Cytometry

Three human cell lines H9 (HTB-176), MDA-MB-231 (HTB-26) and PC3 (CRL-1435) were commercially obtained and cultured as recommended by the supplier (ATCC). Cultured cells were re-suspended in culture media containing Hoescht 33342 (Sigma) and incubated at 37°C for 60 minutes prior to sorting.

Cultured cells were flow-sorted to isolate individual cells from each of the cell lines according to phase (G0/G1, M/G2 and S). Cells were isolated and sorted using the FACSJazz (Becton Dickinson) at 500 events per second using a 100 micron nozzle. Single cells were defined by gating on forward and side scatter area/width. Phase was inferred from Hoescht 3342 DNA-fluorescent dye, then cells were individually deposited and lysed in wells of a 96-well PCR plate containing 3uL of Cells-to-Ct lysis buffer (Life Technologies). The proportion of cells in G0/G1 phases varied from 54% of PC-3 cells to 73% of H9 cells (Supplementary Figure S1).

### Genes Assayed

A set of 333 probes was designed. It contained cell cycle associated genes and provided coverage of the entire cell cycle based on peak expression and periodicity information derived from an integrated database of cell cycle expression profiling experiments [16]. Non-cell cycle associated genes had primary roles in the inflammatory response and included housekeeping controls without a Cyclebase ranking. Genes with a Cyclebase ranking <1000 were placed in the *ranked* set (n = 119) and all other probes were considered part of the *unranked* set (n = 214).

### cDNA Conversion and Multiplexed Target Enrichment (MTE)

After lysis, RNA was converted to cDNA with SuperScript VILO (Life Technologies). Primers for 333 genes were pooled and cDNA was enriched in a multiplexed amplification (MTE) reaction according to the nCounter Single Cell Expression protocol (NanoString). The MTE samples were hybridized overnight at 65°C with an nCounter CodeSet containing probes for all enriched targets (cell cycle related, unrelated genes and controls) and internal controls as recommended by the manufacturer.

### Statistical Analysis

#### Dichotomization and Thresholding

The nCounter Analysis System reports the number of counts of each observed nucleic acid target. We transformed the counts with a shifted log-2 transformation so that *lCount* = log_2_ (count +1). In examining histograms of the transformed data, *lCount*, we found evidence of bi-modality (e.g. Figures 1C, 2). It has been previously observed [2,9,10] in single cell gene expression that genes may appear “off” in a cell, lacking detectable transcript. Thus we hypothesize that in genes with two clusters apparent, the cluster of smaller *lCount* might represent background noise without detectable expression, and the cluster of larger *lCount* might correspond to *bona fide* signal. The distribution of *lCount* in positive controls, which were added at known concentrations, and negative control probes not occurring in human cDNA, additionally supported this hypothesis (Figure S2).

We used an empirical Bayes, model-based clustering procedure to discriminate between signal and noise clusters. Via maximum likelihood estimation, we fitted a Gaussian mixture model to an omnibus of expression in all genes to insure that both signal and noise clusters were initially present. The parameter estimates from the omnibus were then used to form an empirical Bayes estimate of a prior distribution for Bayesian Gaussian mixture models fit to each gene separately. The function *thresholdNanoString* available in *SingleCellAssay* implements our thresholding framework, while mixture models are estimated with the *flowClust* R package [29].

Let 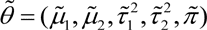 be the MLE estimate of the cluster means, variances, and mixing proportion for the Gaussian mixture model when fit to the omnibus of all genes.

Then for the gene-specific thresholding, the Bayesian formulation of the mixture model imposes the prior for cluster *i* = 1, 2 in gene *g*

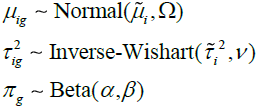

with *μ*_*ig*_,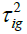 and *π_g_* all mutually independent. The hyperparameters Ω,*ν* were estimated by empirical Bayes using a method-of-moments methods, employing the function *flowClust2Prior* with *kappa = 3, Nt = 5*, while the hyperparameters for the Beta distribution were both set to 5, thus the prior has weight equal to 10 observations in the likelihood. Then for each gene, maximum *a posteriori* (MAP) parameters using Expectation Conditional Maximization, subject to the data for that gene and the prior. Let 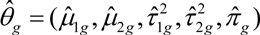 denote the MAP estimate for gene *g.* After ensuring that 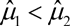 by relabeling the clusters if necessary, we can find the posterior probability that an observation belongs to cluster 1 by considering

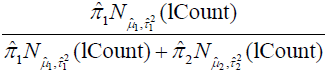

where 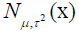 is the Normal PDF with mean *μ* and variance *τ*^2^ evaluated at *x.* Observations with posterior probability >.5 of belonging to the noise cluster were truncated (set to zero) while observations that more likely belonged to the signal cluster were left unchanged.

We denoted the truncated, log-transformed value as the Expression Threshold (*et*) in which the value zero denotes no detectable expression, while positive values correspond to increasing values of log-expression. We model the zero value specially and separately from positive values.

#### Normalization

The log-count measurements were normalized to ensure that the mean signal was comparable across plates. This was done in a stepwise manner. First data were split into three experimental batches, corresponding to cells that were run on different dates, and preliminary signal and noise clusters for each gene were estimated using the above thresholding technique. Then letting lCount_**kip**_ denote the clustered, but un-thresholded log Count for cell *k*, cluster *i* = 1, 2 and plate *p*, we aligned signal and noise clusters by estimating the regression

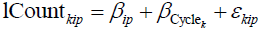

and then subtracting the plate-specific signal and noise intercepts *β*_*ip*_. This aligned the peaks of the signal and noise clusters across plates, akin to a non-pooled version of the ComBat method [30]. The normalized log counts were translated so that the minimum normalized value in each gene was zero. Since no gene was expressed 100% of the time under the preliminary clustering, this was a well-defined procedure. Finally, the normalized data were thresholded jointly to produce the data set used for filtering and testing.

#### Filtering

We adopted a previously published filtering approach [9] based on a robust z-scoring, removing wells with no expression or otherwise outlying in the number of transcripts expressed. We applied the *SingleCellAssay* function *filter* with parameters *nOutlier = 2, SigmaProportion = 2, SigmaContinuous = 5*.

We removed non-variable genes (i.e. detectable expression < 1% in any cell line). 253 of 333 probes passed these filtering criteria and were carried forward in the analysis. After thresholding and filtering we found that the frequency of expression (the rate at which *et* >0 in a gene) varied considerably between genes, with a range of .08-.99 and median value of .72.

#### Amplification Efficiency

We found in examining principle component plots that the first axis of variation corresponded to the number of genes expressed in a well. In cell *k* and gene *g*, let et_*kg*_ be the thresholded log_2_ count, and lCount_*kg*_ be the un-thresholded log_2_ count. Then we defined

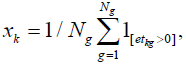

where there are *N*_*g*_ genes total, giving the proportion of genes in the panel that were expressed in a cell. We considered several factors before deciding that *x*_*k*_ corresponded to technical variation that should be removed.

First, higher-order axes of variation corresponded to identifiable biological factors (e.g., phase, cell line). Orthogonality of *x*_*k*_ to the biological axes of variation suggested that this factor was technical in nature[31]. Secondly, many steps in deriving cDNA from live cells could induce technical, cell-specific variation. These steps include incomplete lysis, variation in reverse transcription to generate cDNA and efficiency differences in the multiplexed, amplicon-specific pre-amplification step. Cell-to-cell variability in any of these could appear downstream as a source of variability[32]. Lastly, we have observed similar phenomena in other cDNA-based single cell gene expression experiments, including multiplexed qPCR and single-cell RNA-seq. As *x*_*k*_ contributed variability to our data and appeared to derive from technical rather than biological sources, we chose to adjust for it as a nuisance source of variability.

In fact, *x*_*k*_ is highly correlated to the log-sum of expression

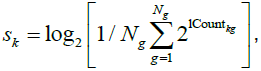

which is equivalent to the log-total read count in RNA sequencing experiments (Supplementary Figure S5). Thus correcting for *x*_*k*_ variability can be seen as a form of normalization, as is typically encountered in RNA-seq.

### Hurdle models for zero-inflated expression

In single cell gene expression, we have previously found that accounting for both changes in the frequency of expression and shifts in the PEM produces more sensitive measures of differential expression compared to using either the frequency or the positive values alone, or compared to t-tests on the zero-inflated values [9,33]. We sought to extend this framework to any model that permits a likelihood ratio test on parameters, *e.g.*, generalized linear or generalized linear mixed models, in order to account for additive cell line effects. Let *et*_*k*_ denote the expression threshold in the *k*th cell (so thus suppressing the gene index). Then we model

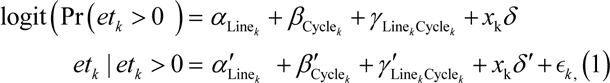

where *α*, *α*′ are cell line effects, *β*, *β*′ are cell cycle effects and *γ*, *γ ′* are interaction effects between cell line and cell cycle, and ο*k* is an independent, normally distributed error. The indices Line_*k*_ and Cycle_*k*_ give the cell line and cell cycle of the *k* th cell. The cell line effects *α*, *α*′ and cell cycle effects *β*, *β*′ are vectors in 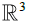 although with the linear constraint that the sum of them is zero, eg, 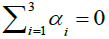, while *γ*, *γ ′* is a matrix in 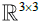 with the constraints that 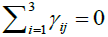 for j = 1, 2,3 and 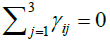 for *i* = 1, 2,3.

The term *x*_k_*δ* accounts for cell-to-cell technical variability resulting from variation in reverse transcription and PCR amplification efficiency (see previous section). Jointly modeling the PCR efficiency along with the biological effects of interest is important as one factor can affect the other. Our modeling framework can be extended to regression-type models when the right hand side is replaced with a general term ***Z****η* for each component, and even to generalized linear mixed models. In general, let *θ* be a vector of parameters for the distribution of 1(*et* > 0) and let *θ*′ be a vector of parameters for *et* | et >0. Then when the distribution of *et* is divided in this fashion, inference about *θ*′ proceeds conditional on *et* >0. The log likelihood is then additive in the *θ* and *θ*′ parameters. Classical hypothesis tests with chi-square asymptotic null distribution, such as Wald or likelihood ratio tests on specific components of *θ* and *θ*′ are null can be conducted separately. Then the test statistics are added together, combining and summarizing the evidence from the two processes, with the degrees of freedom in the null distribution doubled for the purpose of assigning significance. This approach is dubbed the “Hurdle” model and has been used in economics for several decades [34,35].

#### Application of Hurdle Model to tests of Cell Cycle Expression Regulation

For each gene, we test whether the cell cycle effect, (*β*, *β*′), was equal to zero. The log-likelihoods under both models *M*_1_ : (*β*, *β*′) ≠ 0 and *M*_0_ : (*β*, *β*′) = 0 are compared. Let *λ*_0_ and *λ*_1_ be −2 times the log-likelihood under models *M*_0_ and *M*_1_, respectively. Then *λ*_cycle_ = *λ*_0_ − *λ*_1_ gives Wilks’ likelihood ratio statistic, and in large samples, the null hypothesis of no cycle effect can be tested by comparing *λ*_cycle_ to a chi-square distribution with four degrees-of-freedom, as there are three cycles, but with a linear constraint, hence two degrees in each of the discrete and continuous *et* components.

#### Union-Intersection Test of Cell Cycle Expression Regulation

Equivalently, *M*_0_ : (*β*, *β*′) = 0 can be represented as the intersection 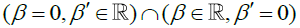. This permits forming a union test of *M*_0_ versus *M*_1_. For a test of size *α* of *M*_0_, the rejection region is then

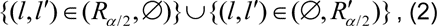

where *l* and *l*′ are the discrete and continuous LRT test with rejection regions *R*_*α*/2_ and 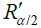, respectively. In other words, this means to reject the intersection hypothesis at level *α* if either discrete or continuous test reject at level *α*/2.

#### Proportion of Deviance Explained by Cell Cycle

In order to calculate the proportion of deviance explained by cell cycle, we compare our Hurdle model given by (1) to the same model where all cell cycle effects are omitted (i.e. (*γ*, *γ′*) = (*β*, *β*′) = 0). Let *λ*_*a*_ be −2 times the log-likelihood under this alternative model. The cell cycle deviance ratio is calculated as *d_cycle_* = (*λ_a_* − *λ*_1_)/*λ_a_*, directly analogous to the calculation of the coefficient of determination *R*^2^ in linear least squares.

The deflated cell cycle deviance ratio is calculated as *d_cycle_* / max (*d_cycle_*), where max (*d_cycle_*) = .27 and occurs in gene TOP2A.

### Network Estimation

We extend the conditional, neighborhood-based algorithm of Meinshausen-Bulmann [36] to estimate co-expression networks using the Hurdle model. The standard Meinshausen-Bulmann algorithm uses L1-penalized regressions to estimate partial correlations between vertices (genes) by treating each vertex as a dependent variable in a regression that includes all other vertices as independent variables. If the vertices are jointly Gaussian, non-zero coefficients correspond to statistical dependences between vertices, conditional on all other factors and so reflect a Gauss-Markov Random Field. Here, since the distribution of expression in single cells is not multivariate Gaussian, edges in our network correspond to conditional correlations (after possible application of the logit link). Although we do not attempt to show consistency of our proposed approach here, we note that Meinshausen-Bulmann-like methods have been shown to be consistent in estimating non-Gaussian graphical models under fairly general conditions [37,38].

Then for the *k*th cell, following equation (1), we divide expression into discrete and continuous components, so fit regressions of the form

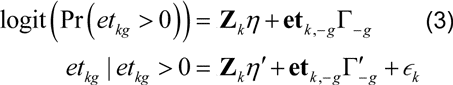

Where *et_kg_* is the expression of the *g*th gene in the *k*th cell, and **et**_*k*,−*g*_ is the expression vector of all except the *g*th gene in the *k*th cell, and **Z*_k_*** is a vector of cellular covariates (eg pre-amplification effect, cell line, cell cycle, and their interaction). We estimate (η,Γ) and (η′,Γ′) separately, with distinct L1 penalties λ and λ′ for Γ and Γ′ using the R package glmnet [39]. Unpenalized vector parameters η and η′ adjust for pre-amplification effect *x*_*k*_; cell line and cell cycle.

#### Combining Networks

We connect genes *g*_1_ and *g*_2_ if any one of 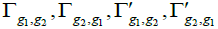 is non-zero at their respective penalties thus take the union of the symmetrized sub-networks. To select the penalty parameters, we fix a number of edges *e*, then find *λ* and *λ*′ (constant across genes) so that *e*/2 edges enter from each of Γ and Γ′. Other ratios of edges are easily attained by choosing *λ* and *λ*′ appropriately.

#### Cellular and Marginal Co-Expression

Even when expression is measured in single cells, co-expression estimates may reflect cluster-specific shifts in mean expression rather than cellular co-expression. Let *G*_1_,*G*_2_ be two genes, and let *Z* be a clustering factor that affects expression of at least one of *G*_1_,*G*_2_. Then an elementary probability calculation shows that

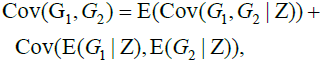

so that the unadjusted estimate of covariance Cov(G_1_, *G*_2_) will include marginal shifts in the means E(*G*_1_ | Z), *E*(*G*_2_ | Z) as well as the average covariance E(Cov(*G*_1_, *G*_2_ | Z)). If *Z* is measured, then it can be used to adjust the regressions in equation (3) to remove the effect of shifts in the mean and so isolate the effect of E(Cov(*G*_1_,*G*_2_ | Z)).

## Acknowledgements

The authors would like to thank Seely Kaufmann and Rich Boykin for bioinformatics support associated with CodeSet design.

## Supplemental Material

**Figure S1.**
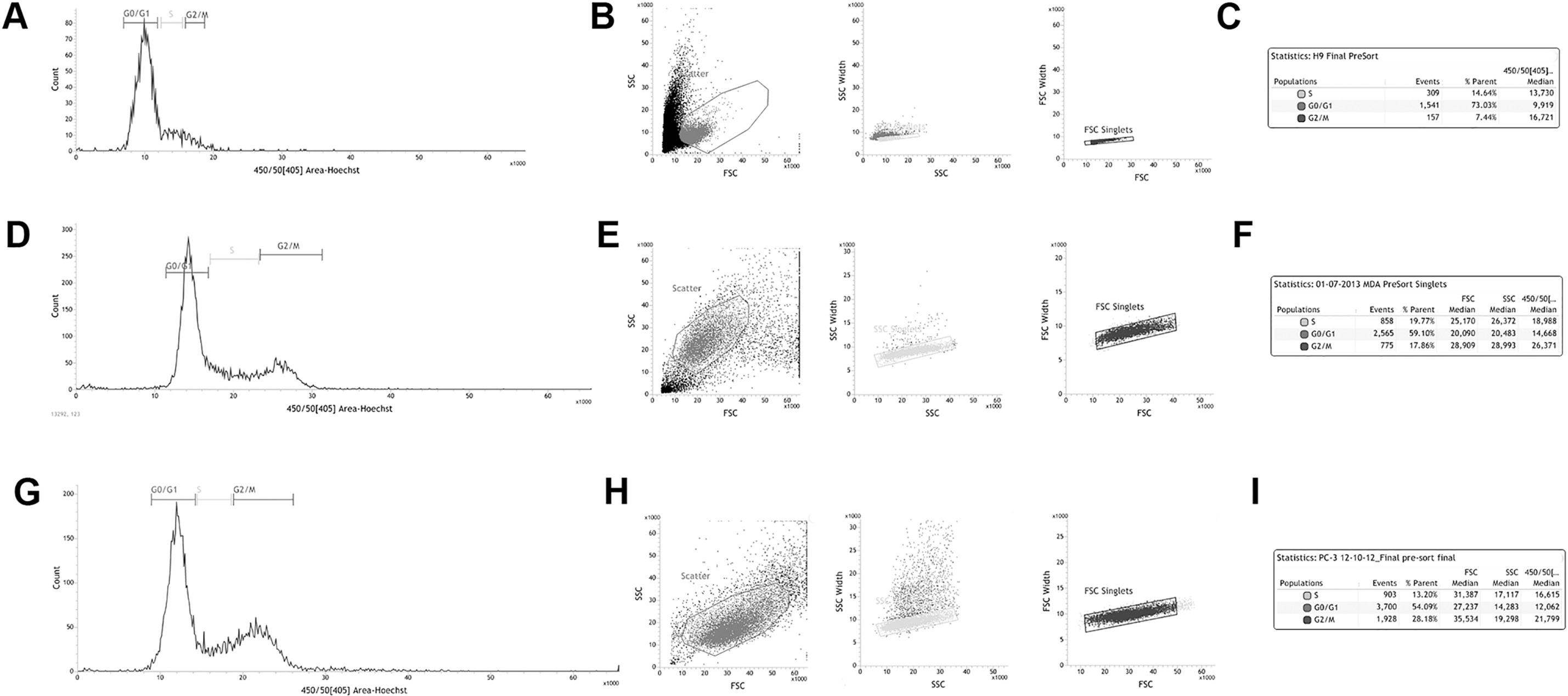
Separation of asynchronously cycling cells into three cell cycle phase populations (G0/G1, S and G2/M) via fluorescence activated cell sorting (FACS). H9 (A), MB-231 (D) and PC3 (G) cells were sorted based on DNA content as determined via retention of Hoechst 33342 dye. Individual cells were gated on based on forward scatter versus side scatter for H9 (B), MB-231 (E) and PC3 (H) populations. The number and percentage of H9 (C), MB-231 (F) and PC3 (I) cells in a given phase within the asynchronous population as determined by FACS analysis.

**Figure S2.**
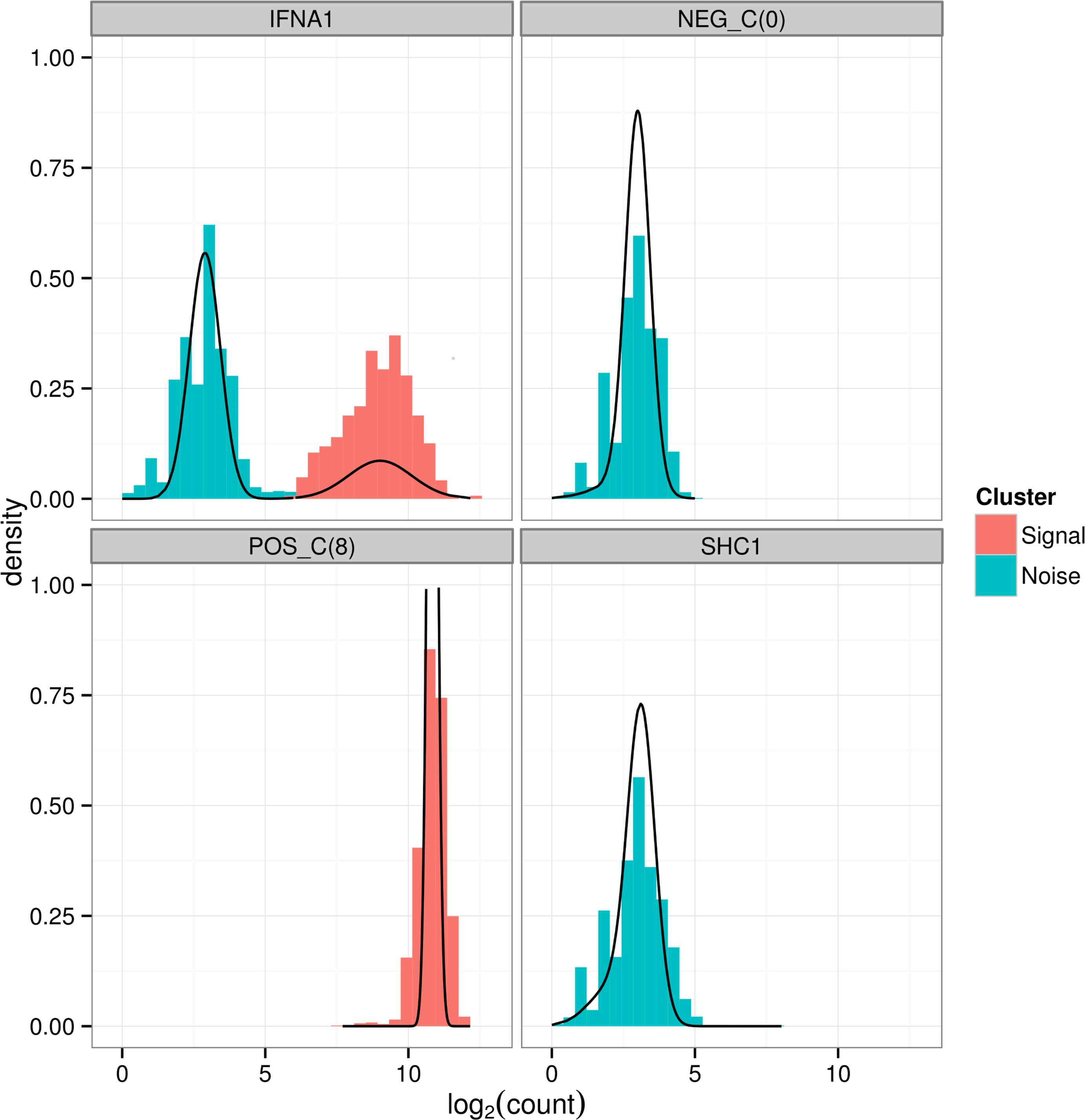
Histogram of log Counts of mRNA for various controls and genes. Positive control primers (for which 100% expression is expected), negative control primers (for which no expression is expected) and two genes with different expression frequencies are shown. The estimated Gaussian mixture densities from the Empirical Bayes model are superimposed.

**Figure S3.**
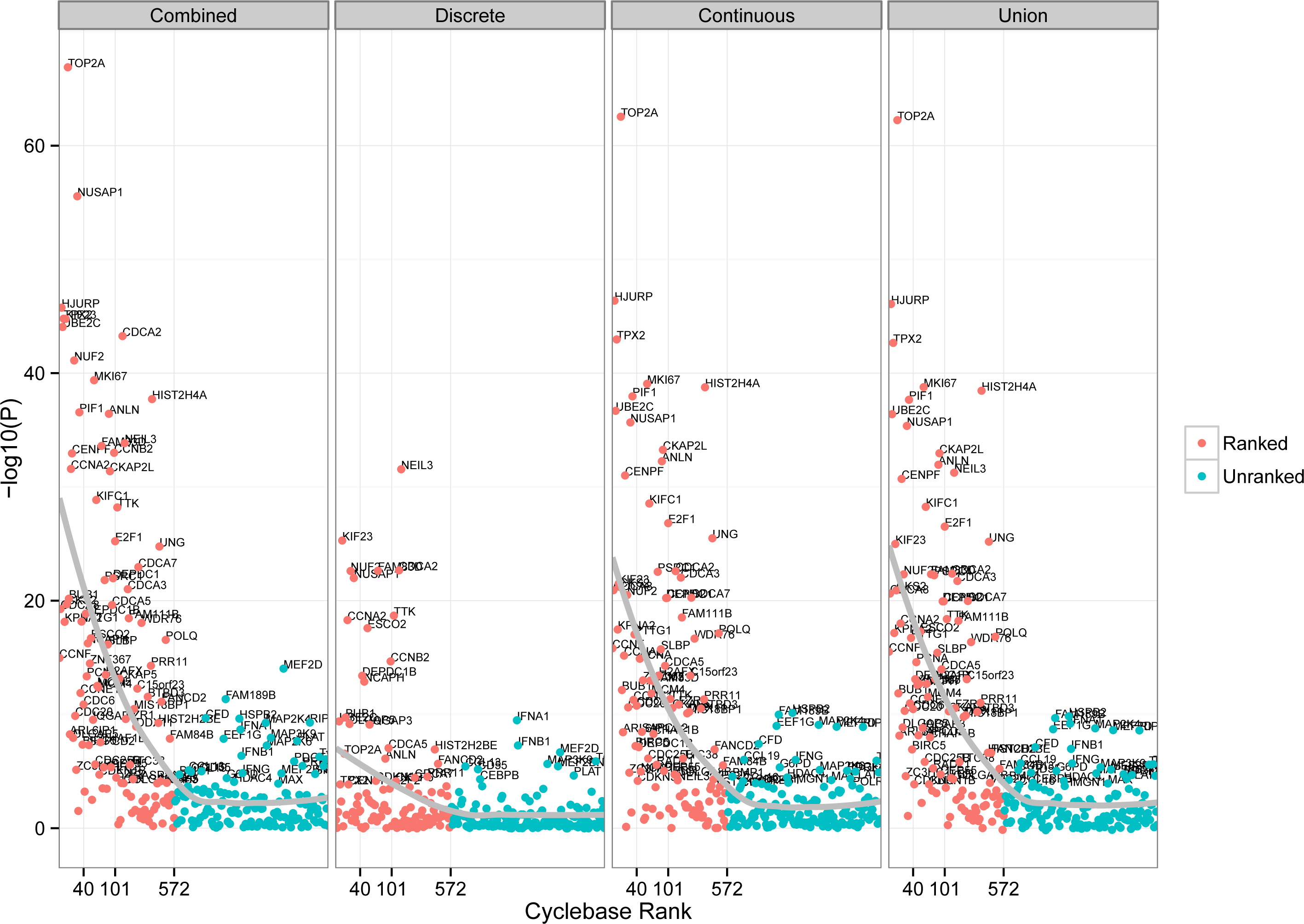
P values testing for differential expression in the Hurdle model decompose into discrete and continuous portions, and a union-intersection on the parameter set. Grey lines indicate average (loess smoothed) P-value for a given gene rank. Both discrete and continuous components offer information about differential expression, and combining them via the Hurdle model offers more sensitive detection of ranked genes compared to a union-intersection test.

**Figure S4:**
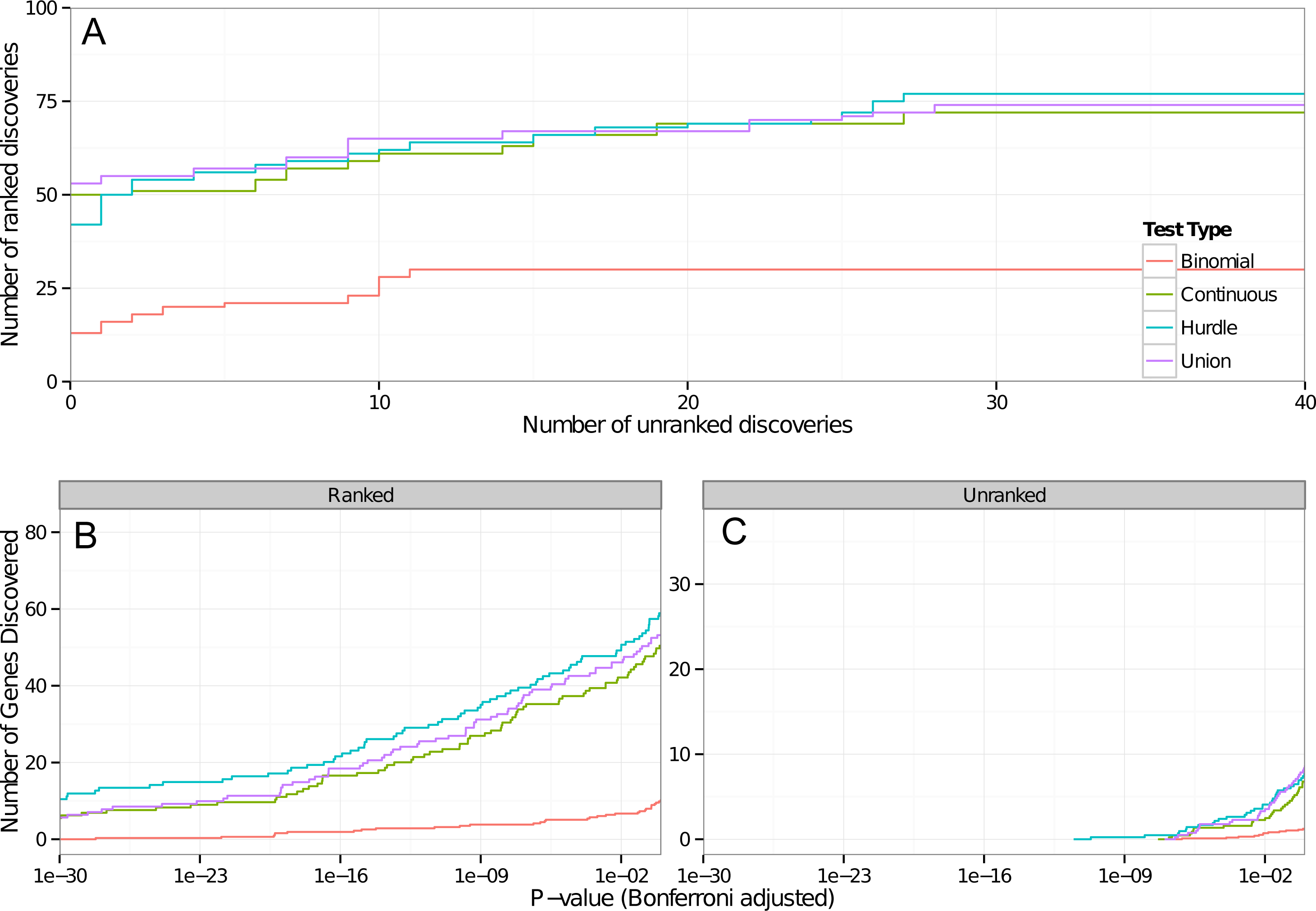
Pseudo ROC plot (A) and number of discoveries versus Bonferroni-adjusted P values for ranked (B) and unranked (C) genes. In panel A the number of discoveries in *ranked* genes is plotted against the number of discoveries in *unranked* genes as the level of the test varies. A discovery in a ranked gene, as it has been previously found to be cell-cycle regulated, is more biologically plausible than a discovery in an unranked gene, so the number discovered at a given level is plausibly related to the sensitivity of a test. Likewise, the number of discoveries in unranked genes may be plausibly related to the specificity of the test. In panels B and C the absolute number of discoveries in ranked and unranked gene sets are plotted for various P-values. In both panels, the binomial model uses logistic regression on dichotomized expression values, while the continuous model uses only values with positive expression. All models adjust for cell line and pre-amplification efficiency. The Hurdle, Union and continuous tests are largely equivalent when judged by their area under the curve of the panel A; however the Hurdle is more sensitive than the continuous or union when judged by absolute number of discoveries in panel B.

**Figure S5:**
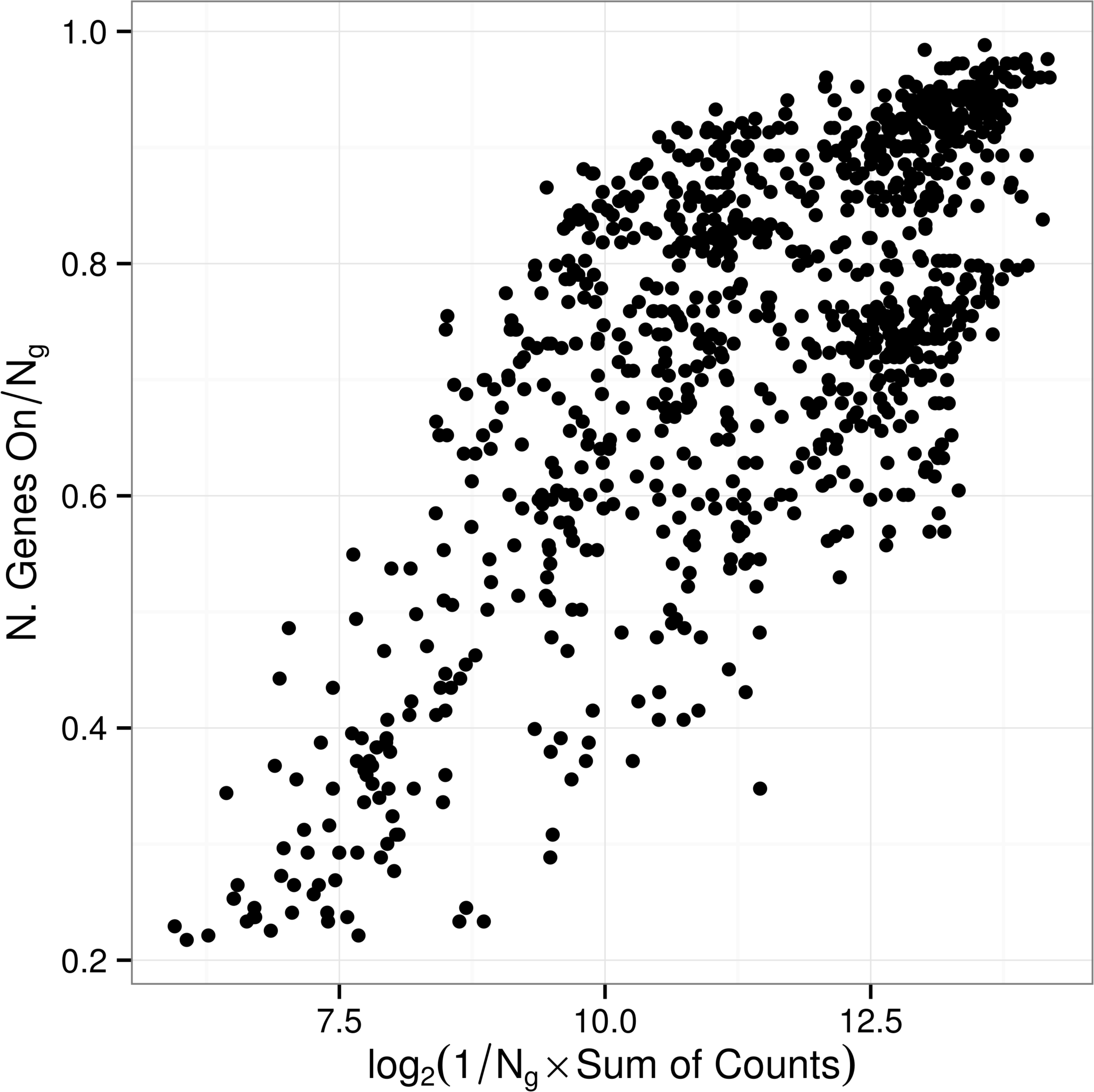
The proportion of expressed genes is related to the log-sum of expression in each cell in our panel of Ng = 253 genes.

**Figure S6:**
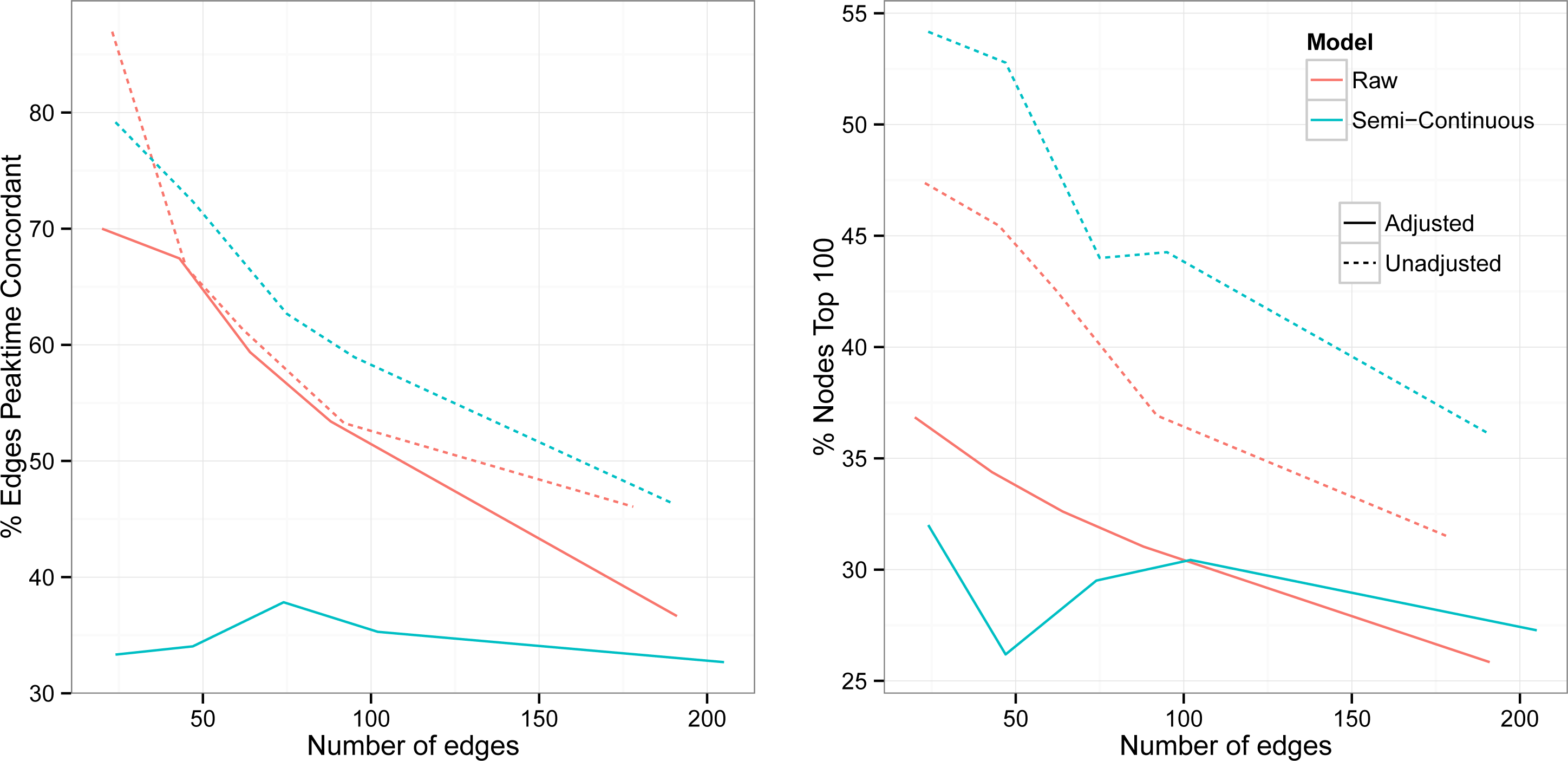
The percent of edges joining nodes with same marginal peak time (A) and percent nodes in top 100 of CycleBase (B) as the number of edges in the network varies from 30-240. The hurdle adjusted for cell cycle selects half as many edges with the same peaktime compared to the adjusted or unadjusted raw models, while the unadjusted hurdle selects modestly more peaktime concordant edges than the raw models, especially for richer (>100 edges) networks. A similar phenomenon occurs when examining the distribution of nodes. The unadjusted hurdle tends to select more nodes with previous description of marginal expression regulation. After adjustment, it selects the fewest nodes out of the four models.

**Figure S7:**
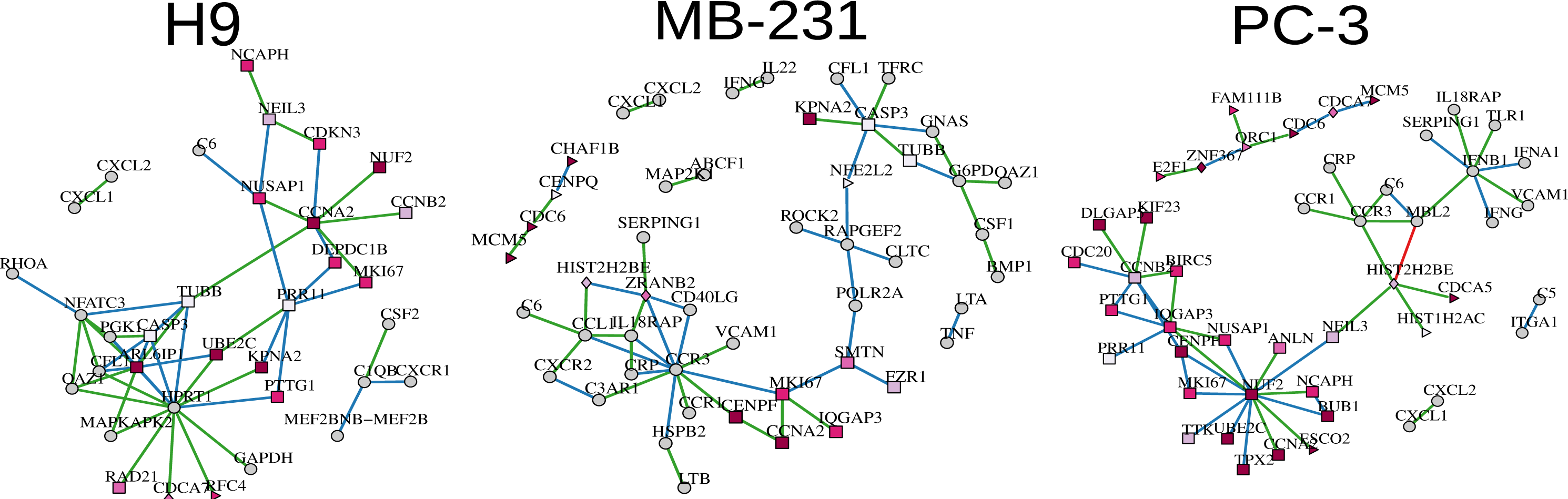
Semi-Continuous Hurdle Model Networks, Adjusted for cell cycle, stratified by cell line.

